# Unconventional Interplay Between GPCRs and RTKs Signaling Pathways Through SH2 Domain-Containing Proteins

**DOI:** 10.64898/2026.04.02.716162

**Authors:** Pedro H. Scarpelli Pereira, Arturo Mancini, Boubacar S. Traore, Hiroyuki Kobayashi, Viktoriya Lukasheva, Christian Le Gouill, Laurent Sabbagh, Michel Bouvier

**Affiliations:** Institute for Research in Immunology and Cancer, and Department of Biochemistry and Molecular Medicine, Université de Montréal, Montréal, QC, Canada; Kainova Therapeutics Canada, Saint-Laurent, QC, Canada

**Keywords:** G protein-coupled receptor, receptor tyrosine kinase, crosstalk, SH2 domain translocation, cell signaling

## Abstract

Crosstalk across two major receptor families involved in signal transduction, namely receptor tyrosine kinases (RTKs) and G protein-coupled receptors (GPCRs), have been observed at different levels of their signaling cascades. Using newly developed enhanced bystander bioluminescence resonance energy transfer (ebBRET)-based biosensors that monitor the recruitment of SH2 domains to activated RTKs, we assessed the ability of GPCRs to modulate cellular localization of SH2 domains. Receptor-mediated activation of either Gα_q/11_ or Gα_12/13_ but not Gα_s_ or Gα_i/o_ (e.g., thromboxane A2 receptor, TPα, and type-2 protease activated receptor, PAR2) resulted in the plasma membrane (PM) dissociation of SH2 domains derived from RTKs effectors such as GRB2, STAT5 and PLCγ1. The role of Gα_q/11_, Gα_12/13_, Rho and downstream kinases in the subcellular SH2 domain redistribution was further confirmed using both pharmacological and genetic approaches. BRET imaging and spectrometric analyses showed that the dissociation of SH2 domains from the PM was accompanied by their accumulation in the nucleus and a reduction in RTK signaling activity, as determined using a STAT5 transcriptional assay. The effect of Gα_q/11_ and Gα_12/13_ activation on STAT5 transcriptional activity was observed both in engineered systems and in HeLa cells endogenously expressing all the components of the regulatory mechanism. The Gα_q/11_ / Gα_12/13_-mediated redistribution of SH2 domain-containing proteins represents an undescribed mechanism through which GPCRs regulate RTKs activity.

**Significance Statement:** This study reveals a novel crosstalk mechanism between G protein coupled receptors and receptor tyrosine kinases showing that Gα_q/11_ and Gα_12/13_ activation triggers Rho-dependent translocation of SH2-containing effector proteins, such as GRB2, PLCγ1 and STAT5. This process causes compartmentalization inside the nucleus and thus reduces their availability at the plasma membrane, leading to attenuated RTK responses.

## Introduction

G protein-coupled receptors (GPCRs) and receptor tyrosine kinases (RTKs) are two major classes of cell surface receptors that regulate critical physiological processes. GPCRs are known to signal mainly through four families of G proteins, each engaging different downstream targets and biological outcomes. While Gα_s_ and Gα_i/o_ families enhance and reduce intracellular cAMP levels, respectively (1), activation of Gα_q/11_ and Gα_12/13_ are associated, among other pathways, to the Rho pathway that is also regulated by RTKs (2, 3).

RTKs undergo homodimerization and cross-phosphorylation on tyrosine residues upon ligand binding. The main mechanism of signal transduction by RTKs is the recruitment of effectors to the phosphotyrosine (phospho-Tyr) residues after receptor activation. These proteins belong to different families, including scaffolding proteins (e.g., GRB2), enzymes (e.g., PLCγ1) and transcriptional factors (e.g., STAT5). The interaction between these accessory proteins and RTKs is mediated by Src Homology 2 (SH2) domains, which specifically recognize phospho-Tyr residues and adjacent amino acids on activated RTKs (4).

While traditionally studied as independent signaling hubs, GPCRs and RTKs pathways crosstalk to enable integrated cellular responses to diverse stimuli. This interplay influences essential biological functions, including cell proliferation, differentiation, metabolism, and motility, with implications in development, homeostasis, and diseases such as cancer and cardiovascular disorders (5, 6). Although the primary mechanisms and machinery involved in RTKs and GPCRs signaling are fundamentally distinct, crosstalk between these two major signaling hubs can greatly broaden the range of cellular responses. For instance, RTKs activation can be mediated by ligands released by matrix metalloproteinases into the extracellular environment as a consequence of GPCRs stimulation (6). Conversely, RTKs may also undergo ligand-independent activation through intracellular kinase cascades triggered by GPCRs, such as those involving Src or phosphatidylinositol-3 kinase (PI3K) (5). The production of reactive oxygen species also represents a mode of transactivation; the Formyl Peptide Receptor 2 and the neurotensin 1 receptor can activate EGFR (7, 8) and TrkA (9) via this mechanism. In addition to transactivation, several points of downstream convergence exist between RTKs and GPCRs signaling pathways, including the activation of Rho, Src, PI3K, ERK1/2, and MAPK (10).

In contrast to the well-documented mechanisms of transactivation, examples and underlying mechanisms of trans inhibition remain scarce. Activation of cannabinoid receptors CB1 and CB2 downregulates EGFR expression, resulting in blunted cellular responses in cancer cells (11, 12). Rigo, A. et al (2012) described that the CXCR4 agonist CXCL12 and its G protein biased variant [N33A]CXCL12 inhibited EGFR phosphorylation in 5637 bladder carcinoma and HeLa cells (13). On the other hand, Tunaru, S. et al (2025) reported inhibition of EGFR signaling by the orphan receptors GPR27 and GPR173 in a G protein and β-arrestin-independent manner when activated by adrenergic ligands, though a direct binding interaction was not confirmed (14). The molecular mechanisms responsible for the complex GPCR-RTK trans inhibition are still poorly understood, and there is a lack of specific tools to directly study the impact of GPCRs stimulation on the mobilization of RTKs downstream effectors.

Using enhanced bystander bioluminescence resonance energy transfer (ebBRET)-based, SH2 domain specific biosensors to quantify RTKs activation, we describe a novel mechanism of trans inhibition between Gα_q/11_- and Gα_12/13_-coupled GPCRs and RTKs. Our data revealed that GPCRs-induced activation of Rho attenuates RTKs signaling by promoting the nuclear localization of effector proteins containing SH2 domains, rendering them unavailable for interaction with RTKs at the plasma membrane (PM).

## Results

### RTKs and GPCRs have opposite effects on SH2 domain-containing proteins

To assess how GPCRs stimulation could affect downstream RTKs signaling, we took advantage of an ebBRET-based RTKs biosensor platform that was developed to quantify RTKs activation and signaling (15). Briefly, the platform consists of i) various *Renilla reniformis* luciferase (rLuc2)-tagged SH2 domains known to interact with RTKs in a phospho-Tyr-dependent manner, and ii) PM-anchored (via the K-Ras CaaX motif) *Renilla reniformis* GFP (rGFP) (Figure 1A). This enables spatiotemporal quantification of SH2 domain recruitment to the PM upon RTK activation without the need to tag the receptor under study. Figure 1B illustrates the effect of EGFR activation on the SH2 domain-based sensors derived from SHP1, SHP2, TNS2, PIK3R2-d2, PIK3R1-d1, PLCγ1, GRB2, and GRB14. EGF stimulation of cells co-expressing EGFR and the different sensors increased BRET signals for all sensors (from 20% and 72% vs. vehicle), consistent with the known action of EGFR on SH2 recruitment and our previous report for a subset of these sensors (15).

**Figure 1.**
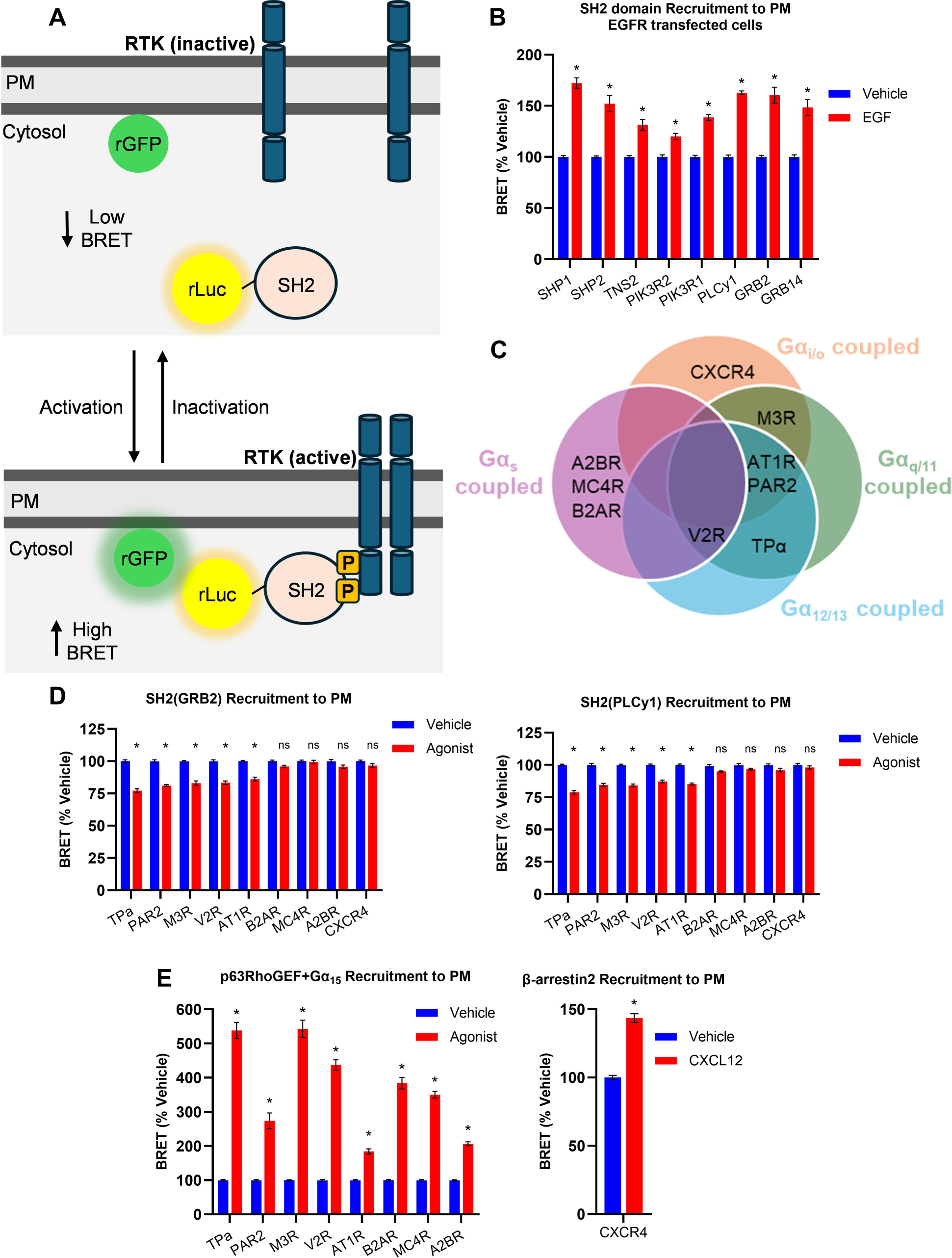
Effect of GPCRs with distinct G protein activation profiles on SH2 biosensor mobilization. (A) Schematic representation of ebBRET-based assay to measure the recruitment of SH2 domains to the plasma membrane (PM). In the basal state, rGFP is anchored to the PM while SH2-rLuc2 is diffusely located throughout the cytoplasm. Upon stimulation and autophosphorylation of an RTK, the SH2-rLuc2 domains are recruited to the phospho-Tyr residue. This increases the concentration of rLuc2 at the PM, the same compartment where rGFP is anchored, thus allowing for efficient resonance energy transfer and resulting in an increase in the BRET signal. (B) HEK293T cells transfected with rGFP-CAAX, EGFR and various SH2 domains fused to rLuc2: SH2(SHP1), SH2(SHP2), SH2(TNS2), SH2(PIK3R2), SH2(PIK3R2), SH2(PLCγ1), SH2(GRB2), and SH2(GRB14) and stimulated with 100nM EGF. (C) Venn diagram representing overlapping G protein coupling profiles of GPCRs used in the tested conditions. HEK293T cells were transfected with rGFP-CAAX, SH2(GRB2)-rLuc2 (D, left), SH2(PLCγ1)-rLuc2 (D, right), and one GPCR coupling to a distinct set of G proteins (TPα: Gα_q/11_ and Gα_12/13_ coupled; B2AR: Gα_s_ and Gα_z/oB_ coupled; PAR2: Gα_q/11_, Gα_i/o_, Gα_12/13_ coupled; MC4R: Gα_s_ coupled; A2BR: Gα_s_ coupled; M3R: Gα_q/11_, Gα_i/o_ and Gα_12/13_ coupled; V2R: Gα_s_, Gα_12/13_, Gα_q/11_ coupled; AT1R: Gα_q/11_, Gα_i/o_, Gα_12/13_ coupled). Since Gα_15_ is a promiscuous G protein that binds well to most tested GPCRs, the p63RhoGEF-rLuc2+Gα_15_ biosensor was used as a receptor activation control (E, left). β-arrestin2 recruitment was used as a control for CXCR4 activation since it does not couple to Gα_15_ (E, right). Before reading, cells were stimulated with a single dose of the agonist corresponding to the GPCR used (TPα: 100nM U46619; B2AR: 100nM Isoproterenol; PAR2: 10μM SLIGKV-NH2; MC4R: 1μM αMSH; A2BR: 1μM adenosine; M3R: 1μM carbamylcholine; V2R: 1μM arginine vasopressin; AT1R: 1μM angiotensin; CXCR4: 100nM). Signals were acquired after 10 minutes for p63RhoGEF-rLuc2 or β-arrestin2 biosensors or 45 minutes for SH2(GRB2)-rLuc2 and SH2(PLCy1)-rLuc2 to calculate BRET values. BRET values were calculated by dividing the intensity of light emitted by rGFP (515 nm) by the intensity of light emitted by rLuc2 (400 nm). Results are expressed as percentage change relative to unstimulated control (mean ± SEM, n=3). * Statistically significant difference by two-way ANOVA with Šidák post-test, p<0,05.

To evaluate the impact of GPCRs activation on the PM recruitment of SH2 domain–containing proteins, we tested SH2(PLCγ1) and SH2(GRB2) biosensors with various GPCRs displaying different G protein coupling preferences (Figure 1C). Stimulation of tromboxane A2 receptor (TPα), protease activated receptor (PAR2), M3 muscarinic acetylcholine receptor (M3R), vasopresin receptor (V2R) and angiotensin II type 1 receptor (AT1R) with their cognate agonists caused the dissociation of SH2(GRB2) and SH2(PLCy1) biosensors (Figure 1D, left and right, respectively), while β2 adrenergic receptor (β2AR), melanocortin receptor 4 (MC4R), adenosine 2B receptor (A2BR) and chemokine receptor type 4 (CXCR4) did not elicit any statistically significant change in SH2(GRB2) and SH2(PLCy1) biosensor activity. The absence of an effect is not due to a lack of activation of these receptors as evidenced by the ability of all receptors (except CXCR4) to activate the promiscuously activated G15 (using the p63-RhoGEF EMTA sensor; (17)) (Figure 1E, left). Of note, G15 is not expressed in HEK293 cells and can be used as a broad GPCRs activation control (Human Protein Atlas; proteinatlas.org and (20)). The activation of CXCR4 was validated by its ability to recruit βarrestin2 (Figure 1E, right). Based on the known G protein coupling preference of these receptors ((17) and Figure 1C), our results suggest that only receptors that engage Gα_q/11_ and/or Gα_12/13_ G protein families promote PM dissociation of SH2(GRB2) and SH2(PLCγ1) biosensors.

To assess the generality of the phenomenon, we investigated the impact of TPα activation on larger panel of SH2-containing proteins (SHP1, SHP2, TNS2, PIK3R1, PIK3R2, PLCγ1, GRB2, GRB14) on their subcellular distribution. TPα is a Gα_q/11_ / Gα_12/13_-coupled GPCR involved in processes such as coagulation, vasoconstriction, and cell migration, with roles in cardiovascular diseases and cancer (21). Consistent with our prior observations, stimulation of TPα expressing cells with the synthetic agonist U46619 resulted in a decrease in the BRET signal for all the SH2 domain considered (Figure 2A). This decrease in BRET is specific to receptor activation as illustrated by the ability of the TPα antagonist SQ29548 to block the U46619-promoted dissociation of SH2(PLCγ1) (Figure 2B).

**Figure 2.**
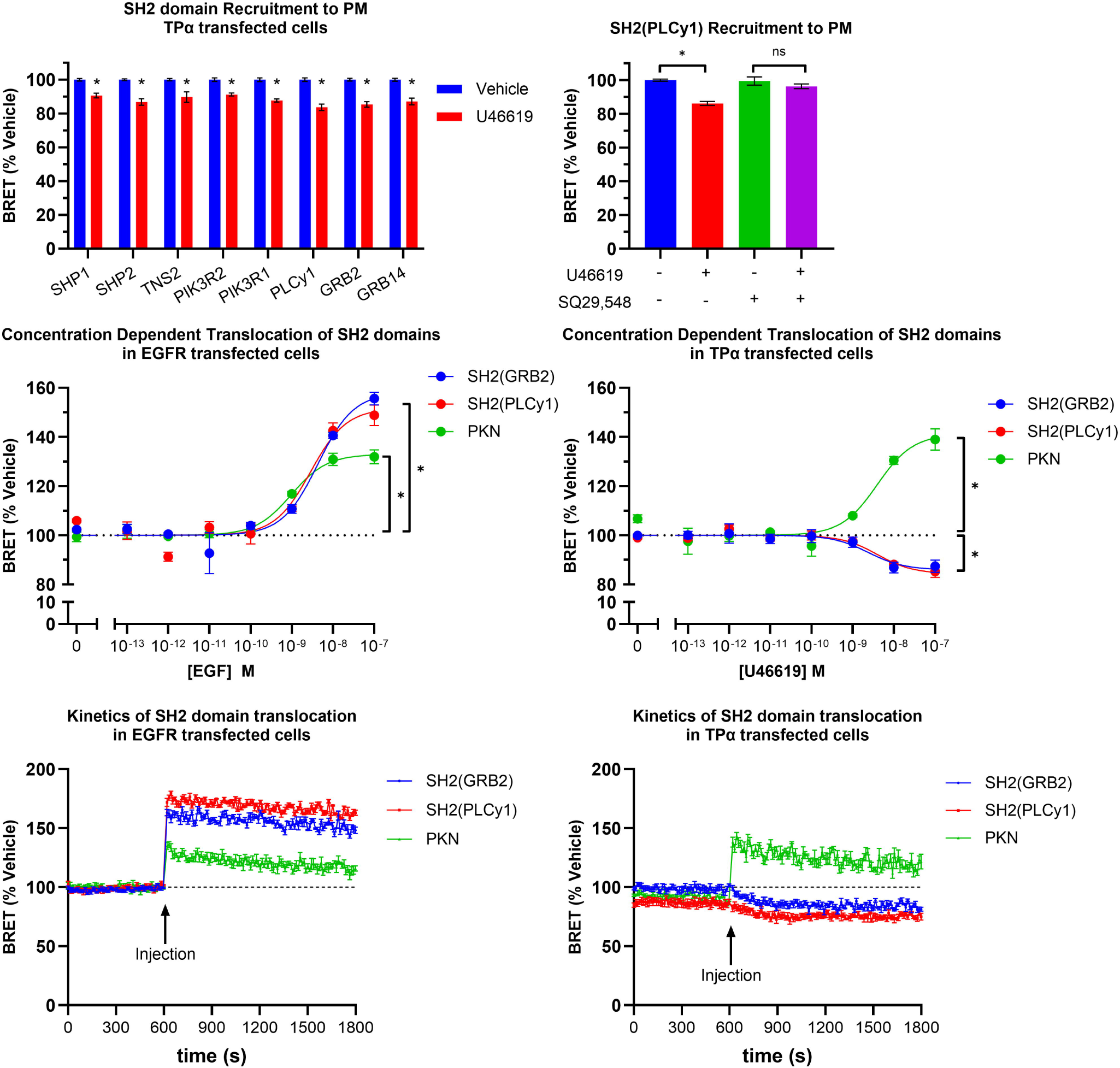
Effect of TPα or EGFR stimulation on the recruitment of SH2 domain biosensors to the plasma membrane (PM). (A) HEK293T cells transfected with rGFP-CAAX, TPα and various SH2 domains fused to rLuc2: SH2(SHP1), SH2(SHP2), SH2(TNS2), SH2(PIK3R2), SH2(PIK3R2), SH2(PLCγ1), SH2(GRB2), and SH2(GRB14) and stimulated with 100nM U46619. (B) SH2(PLCγ1) translocation in the presence of TPα antagonist SQ29548. (C) EGF and U46619 concentration response curves in EGFR (C, left) or TPα (C, right) transfected HEK293T cells using biosensors for measuring the recruitment of SH2(GRB2), SH2(PLCγ1) and PKN (receptor activation control). (D) Kinetics of SH2(PLCγ1) (red), SH2(GRB2) (blue) and PKN (green) translocation to the PM over time in cells transfected with EGFR and stimulated with 10nM EGF (left) or transfected with TPα (right) and stimulated with 100nM U46619. Basal BRET was read for 10 min before addition of agonist. BRET values were calculated by dividing the intensity of light emitted by rGFP (515 nm) by the intensity of light emitted by rLuc2 (400 nm). Values were acquired after 10 min of EGF treatment or after 45 min of U46619 treatment, and results are expressed as percentage change relative to control (mean ± SEM, n=3). * Statistically significant difference by unpaired T-test, p<0,05.

Because the SH2 domains from PLCγ1 and GRB2 yielded the highest signal-to-noise ratio, we proceeded to use these two sensors for a more in-depth characterization of this phenomenon. Further, since PKN is a downstream effector of both EGF- and TPα-promoted Rho activation, it was used as a control to compare each receptor’s activity on SH2 subcellular distribution vs their activity toward their common effector PKN. For this purpose, an ebBRET-based biosensor monitoring the recruitment of PKN to the PM (16) was used. The EGFR-promoted PM recruitment and TPα-promoted PM dissociation of the SH2 domains were found to be time- and ligand concentration-dependent. EGF stimulation promoted the recruitment of SH2(GRB2), SH2(PLCγ1), and the PKN sensors with similar potencies (EC₅₀ = 4.3 nM, 2.8 nM, and 9.6 nM, respectively; Figure 2C, left). Similarly, the potency of the TPα agonist U46619 to induce PKN recruitment (EC₅₀ = 8.4 nM) was similar to its potency in promoting the dissociation of the GRB2 and PLCγ1 SH2 domains (EC₅₀ = 8.6 nM and 8.4 nM, respectively; Figure 2C, right). The similar potency toward PKN recruitment and SH2 dissociation is consistent with the effect on the SH2 distribution being a specific action of the receptor.

In contrast to the similar potencies observed for EGFR and TPα in promoting PM recruitment and dissociation of the SH2 domains, respectively, the kinetics of these responses was different (Figure 2D). These kinetics were compared to that of PKN activation by the two receptors. The BRET signal peaked within 12 seconds of EGF stimulation for the PLCγ1(SH2), GRB2(SH2) recruitment and PKN sensors and gradually declined over the following minutes (Figure 2D, left). Conversely, TPα activation with U46619 resulted in a rapid PKN recruitment (peak response within 12 seconds following ligand addition) whereas the dissociation of PLCγ1(SH2) and GRB2(SH2) was much slower, decreasing gradually over the first 5 minutes post-stimulation and stabilizing thereafter (Figure 2D, right). Such kinetic differences suggest that distinct mechanisms are responsible for the EGFR- and TPα-promoted SH2 domains’ translocation.

### Gα_q/11_ or Gα_12/13_ activation is sufficient and necessary for GPCRs-promoted SH2 dissociation

To directly assess the importance of Gα_q/11_ and Gα_12/13_ signaling in regulating GPCRs-mediated PM dissociation of SH2 domain-containing proteins, we used Gα protein knockout (KO) cells transiently expressing TPα and either the SH2(GRB2), SH2(PLCγ1) or the PKN biosensor. As expected, the activation of the PKN biosensor by U46619-stimulated TPα was completely abolished in Gα protein KO cells but was restored upon re-expression of Gα_q/11_ family members (i.e., Gα_q_, Gα_11_, Gα_14_ and Gα_15_), Gα_12_ or Gα_13_ (Figure 3A). Similarly, the decrease in BRET signals observed for either SH2(GRB2) or SH2(PLCγ1) sensors in WT cells was lost in the Gα protein KO cells but was restored when re-expressing either Gα_q/11_ or Gα_12/13_ family members (Figures 3B and 3C), indicating that the expression of Gα_q/11_ or Gα_12/13_ family members was sufficient to promote SH2 domain dissociation upon TPα activation. To further confirm these observations, we assessed SH2(PLCγ1) biosensor dissociation following PAR2 activation. PAR2 is known to activate Gα_q/11_, Gα_12/13_ and Gα_i/o_ family members (17). Using Gα protein KO cells transfected with PAR2, we observed that rescuing the expression of Gα_q_ or Gα_13_ was sufficient to restore the PAR2-promoted dissociation of SH2(PLCγ1). In contrast, rescuing the expression of Gα_i2_ did not restore the receptor-promoted dissociation of the SH2 domain (Supplementary figure S1). Collectively, these data demonstrate that activation of Gα_q/11_ or Gα_12/13_, but not Gα_i/o_, is sufficient and necessary for SH2 biosensor dissociation from the PM.

**Figure 3.**
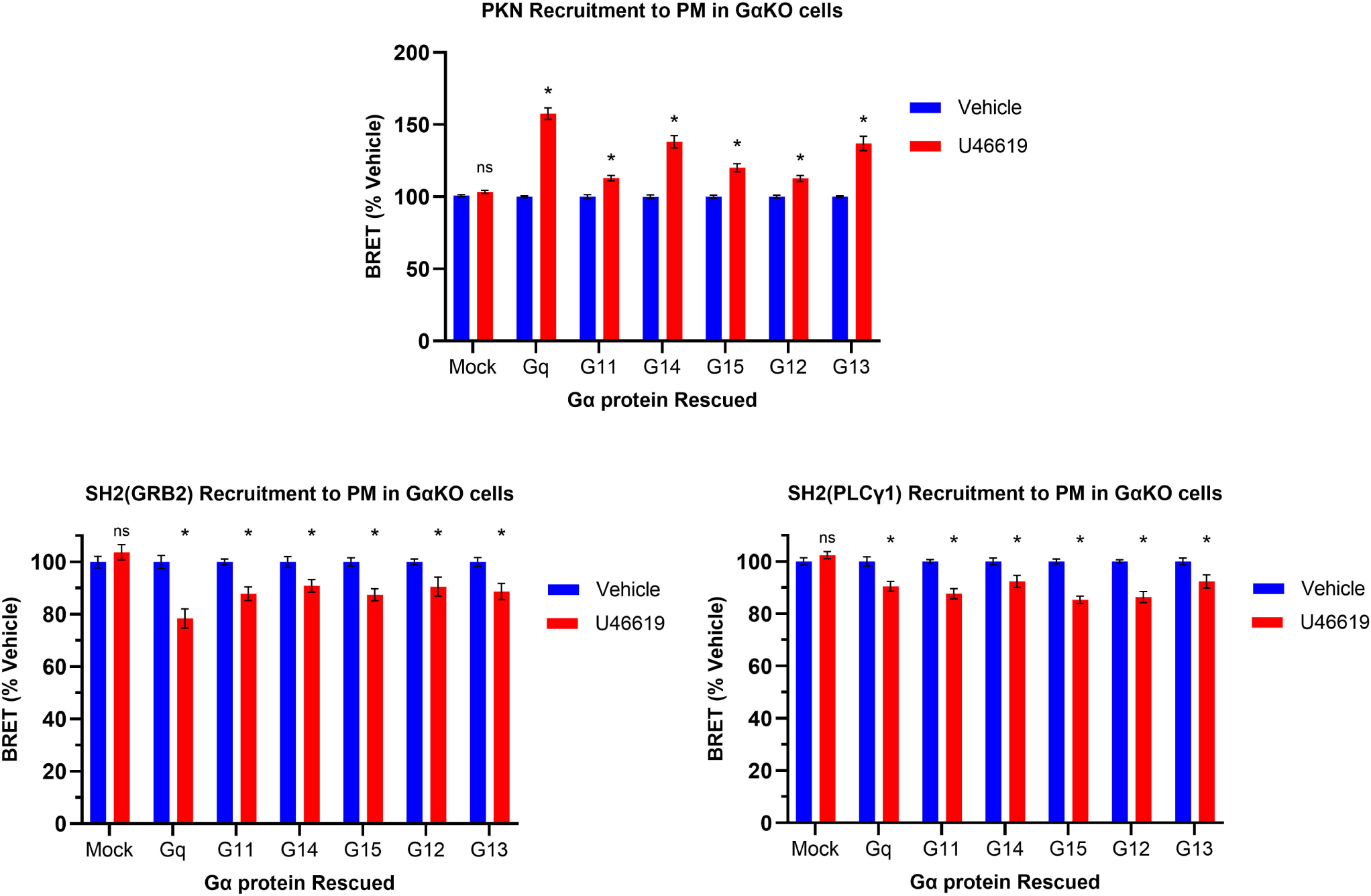
Gα protein dependent SH2(PLCy1) and SH2(GRB2) mobilization by TPα. HEK293T Gα knockout cells were transfected with rGFP-CAAX, TPα and either PKN-rLuc2 (A), SH2(GRB2)-rLuc2 (B) or SH2(PLCγ1)-rLuc2 (C). Each condition was complemented by transfecting 20ng of empty pcDNA3.1 (mock) or one Gα_q/11_ / Gα_12/13_ family member. Cells were treated with 100nM U46619 (red bars) or vehicle (blue bars). Signals were read after 10 minutes for PKN-rLuc2 or 45 minutes of SH2(GRB2)-rLuc2 and SH2(PLCy1)-rLuc2 to calculate BRET values. BRET values were calculated by dividing the intensity of light emitted by rGFP (515 nm) by the intensity of light emitted by rLuc2 (400 nm). Results are expressed as percentage change relative to unstimulated control (mean ± SEM, n=3). * Statistically significant difference by two-way ANOVA with Šidák post-test, p<0,05.

### Gα_q/11_- and Gα_12/13_-promoted SH2 domain translocation is mediated by Rho and downstream kinases

Signaling via Gα_q/11_ and Gα_12/13_ families is known to converge on the activation of Rho, a small GTPase molecular switch involved in multiple cell functions (22, 23). Active Rho transmits signals through several downstream kinases, including STE20-like kinase (SLK/LOK) and Rho-associated kinase (ROCK) (24, 25). To assess the potential role of this pathway in the TPα-promoted SH2 mobilization, we tested the effect of the Rho (CT04) as well as SLK/LOK (Cmpd31) and ROCK (Y-27632) inhibitors. As shown in Figure 4A, CT04 treatment decreased SH2(GRB2) and SH2(PLCγ1) biosensor PM dissociation after stimulation with U46619, indicating that Rho activation plays an essential role in this response. Similarly, inhibition of SLK/LOK with Cmpd31 (Figure 4B) and ROCK inhibition with Y-27632 (Figure 4C) partially reduced the SH2(GRB2) and SH2(PLCγ1) dissociation from the PM. Taken together, these data support an important role for Rho and its downstream kinases in the Gα_q/11_- and Gα_12/13_-mediated SH2 mobilization upon TPα stimulation.

**Figure 4.**
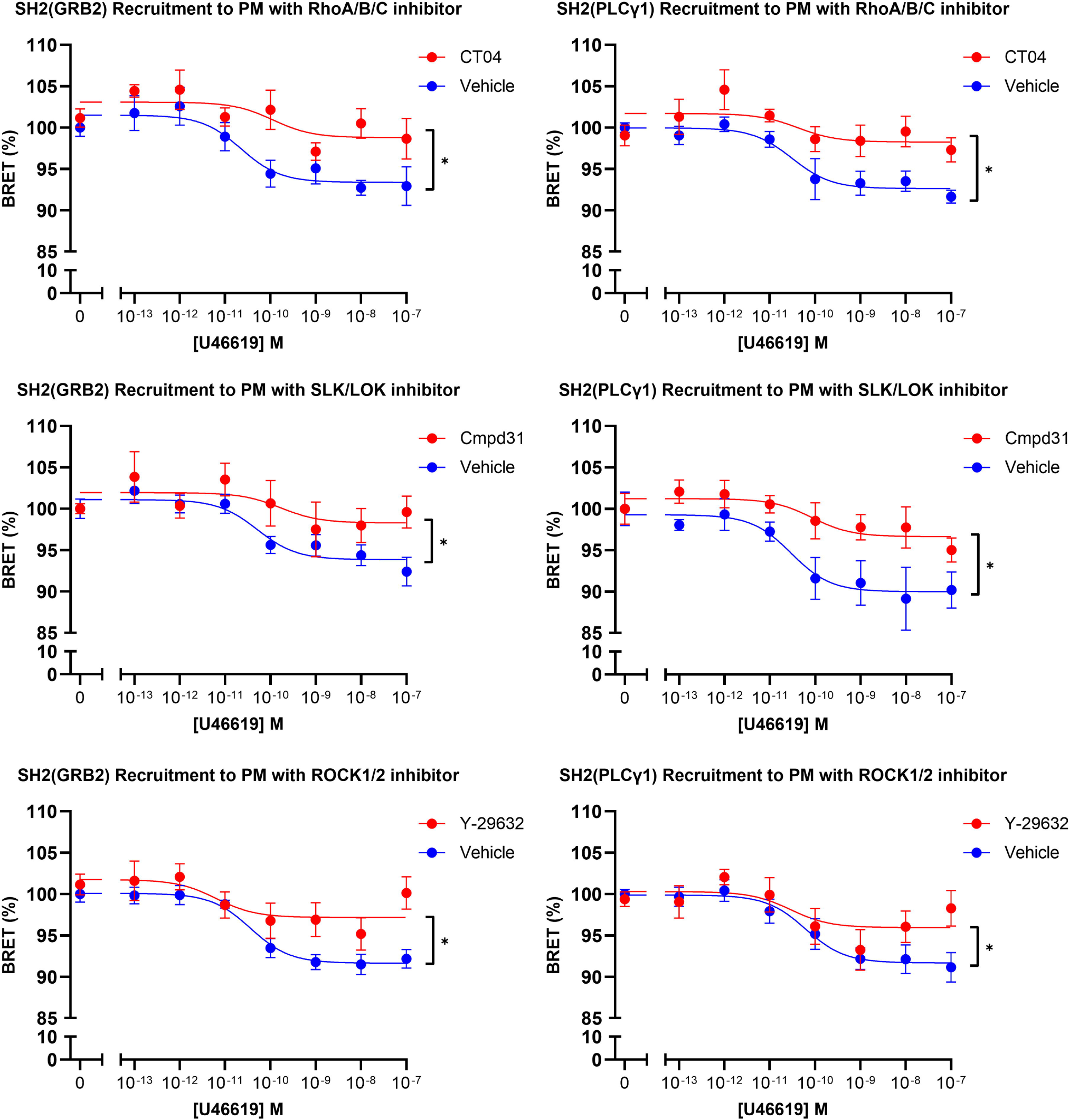
RhoA pathway influences SH2(PLCy1) and SH2(GRB2) mobilization from the plasma membrane. HEK293T cells were transfected with SH2(PLCγ1)-rLuc2 (right) or SH2(PLCγ1)-rLuc2 (left), rGFP-CAAX and TPα. Cells were pre-incubated with 2μg/mL CT04 (RhoA/B/C inhibitor) for 6 hours (A), 1μM Cmpd31 (SLK/LOK inhibitor) for 10 minutes (B), or 10μM Y-29632 (ROCK inhibitor) for 6 hours (C). Cells were then stimulated with a range of 8 concentrations from 0 to 100nM of U46619. Signals were read after 45 minutes to calculate BRET values. BRET values were calculated by dividing the intensity of light emitted by rGFP (515 nm) by the intensity of light emitted by rLuc2 (400 nm). * Statistically significant difference by two-way ANOVA with Šidák post-test, p<0,05.

### SH2 domain dissociation occurs independently of phospho-Tyr

SH2 domains act as specialized molecular adapters that recognize and bind to phospho-Tyr residues within specific sequence motifs on target proteins, including RTKs. We therefore investigated whether the phospho-Tyr binding activity of SH2 proteins is critical for their PM dissociation following GPCRs activation. To this end, we constructed phospho-Tyr binding-deficient mutants of SH2(GRB2) and SH2(PLCγ1) sensors by replacing the arginine of their SH2 phospho-Tyr recognition sites (26, 27) by an alanine (i.e., SH2(GRB2)-R32A or SH2(PLCγ1)-R37A) (Figure 5A).

**Figure 5.**
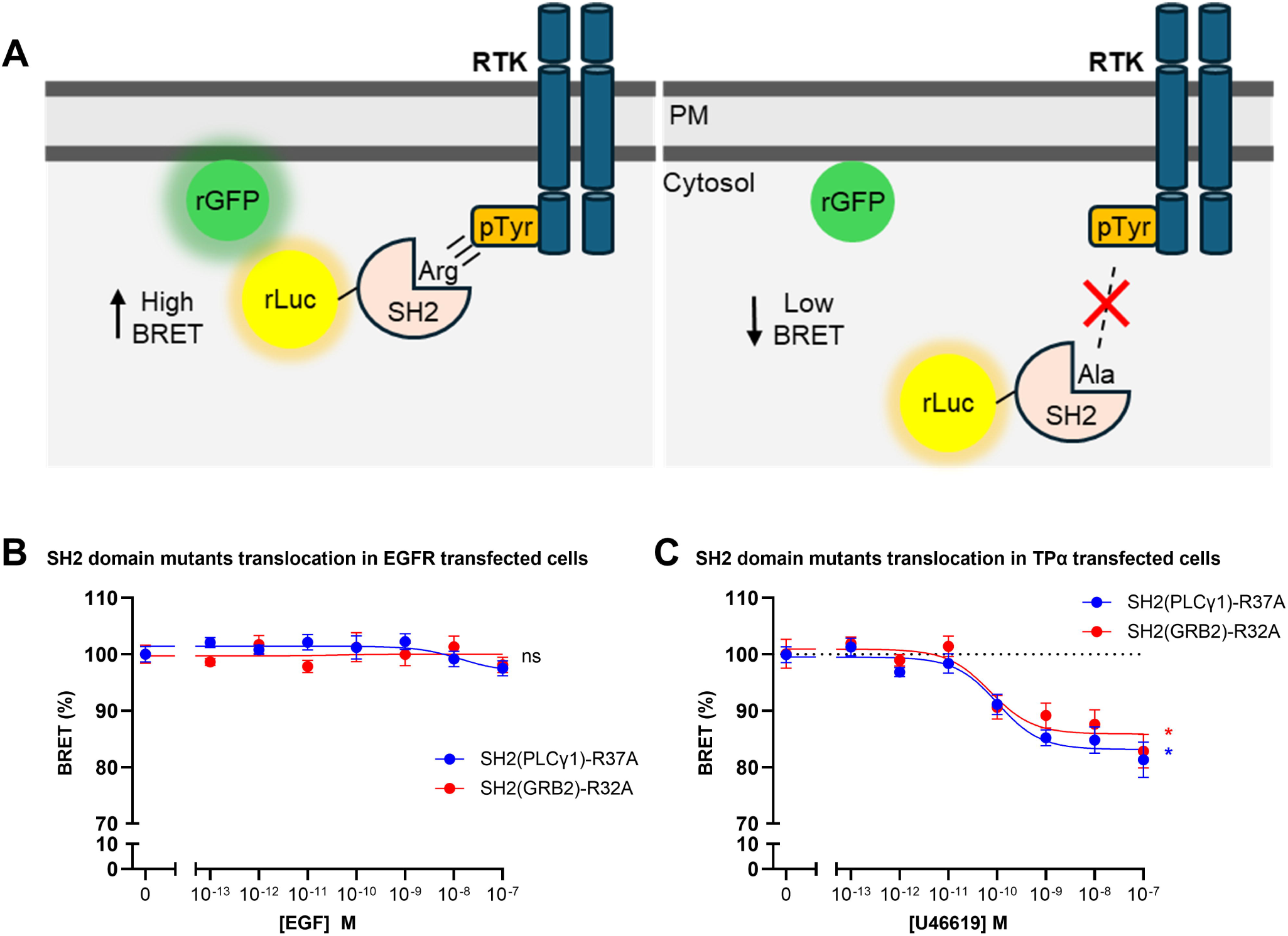
SH2(PLCγ1)-R37A and SH2(GRB2)-R32A mutants unable to bind phosphor-Tyr can still be mobilized following GPCRs stimulation. (A) Schematic representation of the recruitment of SH2 domain sensors by RTKs. SH2 from wild type proteins are recruited to the RTKs due to an interaction between RTKs phosphor-Tyr residues and an arginine (Arg, R) in the SH2 domain, causing an increase in BRET. On the other hand, an Arg (R) / alanine (Ala, A) mutant of the SH2 domain is unable to bind phosphor-Tyr residues, and consequently is not recruited to the plasma membrane, maintaining the BRET values unchanged. (B) HEK293T cells were transfected with rGFP-CAAX, EGFR and SH2(PLCγ1)-rLuc2-R37A or SH2(PLCγ1)-rLuc2-R32A. Cells were then stimulated with 8 concentrations ranging from 0 to 100nM of EGF. (C) HEK293T cells were transfected with rGFP-CAAX, TPα and SH2(PLCγ1)-rLuc2-R37A or SH2(PLCγ1)-rLuc2-R32A. Cells were then stimulated with 8 concentrations ranging from 0 to 100nM of U46619. Signals from U46619 stimulated cells were acquired after 45 minutes, while signals from EGF stimulated cells were acquired after 10 minutes. Concentration response curves were plotted using non-linear fit (three parameters) from three independent experiments (mean ± SEM, n=3).

In contrast to the robust, EGF-concentration-dependent recruitment of WT SH2(GRB2) and SH2(PLCγ1) following EGFR stimulation, no EGF-dependent recruitment of SH2(GRB2)-R32A or SH2(PLCγ1)-R37A was observed (Figure 5B), indicating that these mutations abolished biosensor recruitment to the RTK. On the other hand, TPα stimulation with U46619 promoted a decrease in BRET for the SH2(GRB2)-R32A and SH2(PLCγ1)-R37A biosensors similar to the one observed with WT sensors (Figure 5C). Therefore, the dissociation of SH2 domain-containing proteins occurs independently of the canonical phospho-Tyr recognition site.

### GPCRs promotes translocation of SH2 domain-containing protein to the nucleus

Taking advantage of BRET imaging (18), we tracked the subcellular redistribution of the SH2 domain biosensors following TPα stimulation. As shown in Figure 6A, stimulation with U46619 lead to a time-dependent decrease in BRET between SH2(GRB2)-rLuc and rGFP-CAAX with a concomitant increase in BRET between SH2(GRB2)-rLuc and a nuclear rGFP (i.e., NLS-rGFP). It should be noted that even at the basal level, some BRET is observed between SH2(GRB2)-rLuc and NLS-rGFP indicating that, even before stimulation, some SH2(GRB2) is already present in the nucleus, consistent with the previous observations that GRB2 is partitioned between the cytoplasm and the nucleus (28). We further confirmed the microscopy results via spectroscopic BRET measurements using SH2(GRB2)- and SH2(PLCy1)-rLuc fusions with rGFP targeted to either the PM or nucleus. Activation of TPα with U46619 resulted in a concentration-dependent signal decrease at the PM and increase at the nucleus for both SH2(PLCy1) and SH2(GRB2) sensors (Figure 6B and 6C respectively). Since our SH2 biosensor constructs do not present any nuclear import signal, we tested if the association with nuclear importins would mediate their translocation. Cells treated with the specific importin-β inhibitor Importazole neutralized the change in BRET after TPα stimulation (Figure 6D). Collectively, these data suggest that stimulation of Gα_q/11_- and Gα_12/13_-coupled GPCRs, such as TPα, results in a redistribution of SH2 domain-containing proteins from the PM to the nucleus.

**Figure 6.**
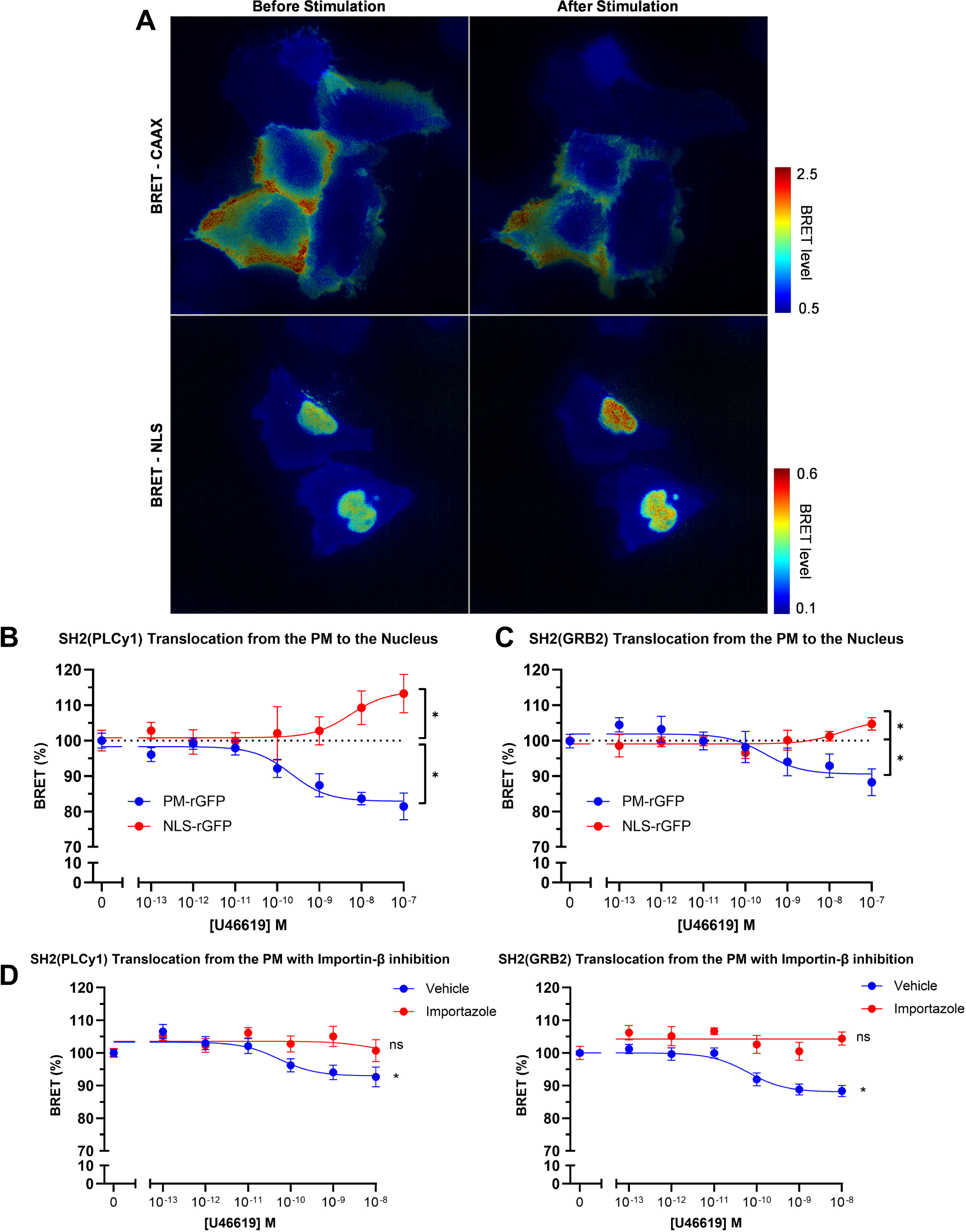
Visualization of (SH2)GRB2-rLuc2 nuclear translocation by ebBRET microscopy. ebBRET imaging was used to monitor nuclear translocation of SH2(Grb2)-rLuc2 in U2OS cells transfected with TPα, rGFP-CAAX or NLS-rGFP and. BRET images were obtained in the presence of 10μM Prolume Purple every 30 seconds for 1 hour after stimulation with 100nM of U46619. Left images are from the same field of view right before stimulation and right column images after 30 minutes stimulation. BRET levels correspond to the ratio of acceptor (rGFP) photon counts divided by donor (rLuc2) photon counts calculated for each pixel. BRET levels are expressed as a color-coded heat map with the lowest being blue, and highest red. Concentration response curves of U46619 in HEK293T cells transfected with TPα, NLS-rGFP or rGFP-CAAX and SH2(PLCγ1)-rLuc2 or SH2(GRB2)-rLuc2 (C). Concentration response curves of U46619 in HEK293T cells transfected with TPα, rGFP-CAAX and SH2(PLCγ1)-rLuc2 (D, left) or SH2(GRB2)-rLuc2 (D, right) in the presence of 40μM importazole (Importin-β inhibitor). Concentration response curves were plotted using non-linear fit (three parameters) from three independent experiments (mean ± SEM, n=3).

### GPCRs attenuates RTK-dependent STAT5 transcriptional activity

Since Gα_q/11_- and Gα_12/13_-coupled GPCRs cause SH2 domain-containing proteins to translocate to the nucleus, we explored the biological significance of this effect by investigating if RTKs-promoted transcriptional response is affected by the lack of available SH2 proteins able to bind activated RTKs at the PM. To test this hypothesis, we used a luminescent transcriptional biosensor reporting STAT5 activation. This reporter consists of a plasmid encoding nanoluciferase (NanoLuc) under a STAT5-specific promoter (see methods and Supplementary figure S2). STAT5 is recruited to phospho-Tyr residues on activated RTKs, undergoing phosphorylation and homodimerization. Phospho-STAT5 then translocates to the nucleus, where it binds the promoter and drives NanoLuc transcription, emitting a luminescent signal that is proportional to the magnitude of the transcriptional response (Figure 7A). We chose STAT5 as a readout because, unlike many other transcription factors, it responds primarily to RTKs and not GPCRs (29), thus avoiding the introduction of confounding GPCR-mediated signals in our crosstalk studies. Indeed, as shown in Figure 7B (left), EGFR stimulated luminescence emission, reflecting STAT5 activity, whereas TPα-stimulation alone did not produce any signal. In addition, the SH2 domain of STAT5 behaves like SH2(PLCy1) and SH2(GRB2), dissociating from the PM following stimulation of TPα (Figure 7B, right). In cells co-transfected with EGFR and TPα, stimulation with U46619 led to a 60% reduction of the STAT5-promoted NanoLuc transcription stimulated by EGF (Figure 7B, red line), indicating that SH2 redistribution prevented the activation of STAT5 by EGFR. In contrast, stimulation of MC4R, a Gα_s_-coupled GPCR, caused no alteration in EGFR-mediated STAT5 signaling (Figure 7C). Pharmacological inhibition of Gα_q/11_ using YM254890 partially reversed U46619’s inhibitory effect on STAT5 activation by EGFR, highlighting the importance of TPα-mediated Gα_q/11_ activation in EGFR STAT5 signaling attenuation (Figure 7D). In addition, TPα stimulation did not affect EGFR promoted STAT5 activation in cells lacking Gα_12/13_ subunits and treated with YM254890 (Figure 7E). Thus, as observed with SH2 domain-containing protein sensors, STAT5 signal attenuation triggered by RTKs activation depends on Gα_q/11_ and/or Gα_12/13_ signaling. TPα stimulation also caused a reduction in platelet-derived growth factor receptor β (PDGFRβ)-induced STAT5 signaling by 50%, indicating that this effect is not exclusive to EGFR (Figure 7F). Finally, experiments in HeLa cells expressing endogenous TPα and EGFR showed that U46619 stimulation also attenuated EGF-induced STAT5 activation (Figure 7G), confirming that the phenomenon is observed with natively expressed receptors and strengthening the physiological significance of this novel crosstalk regulatory mechanism.

**Figure 7.**
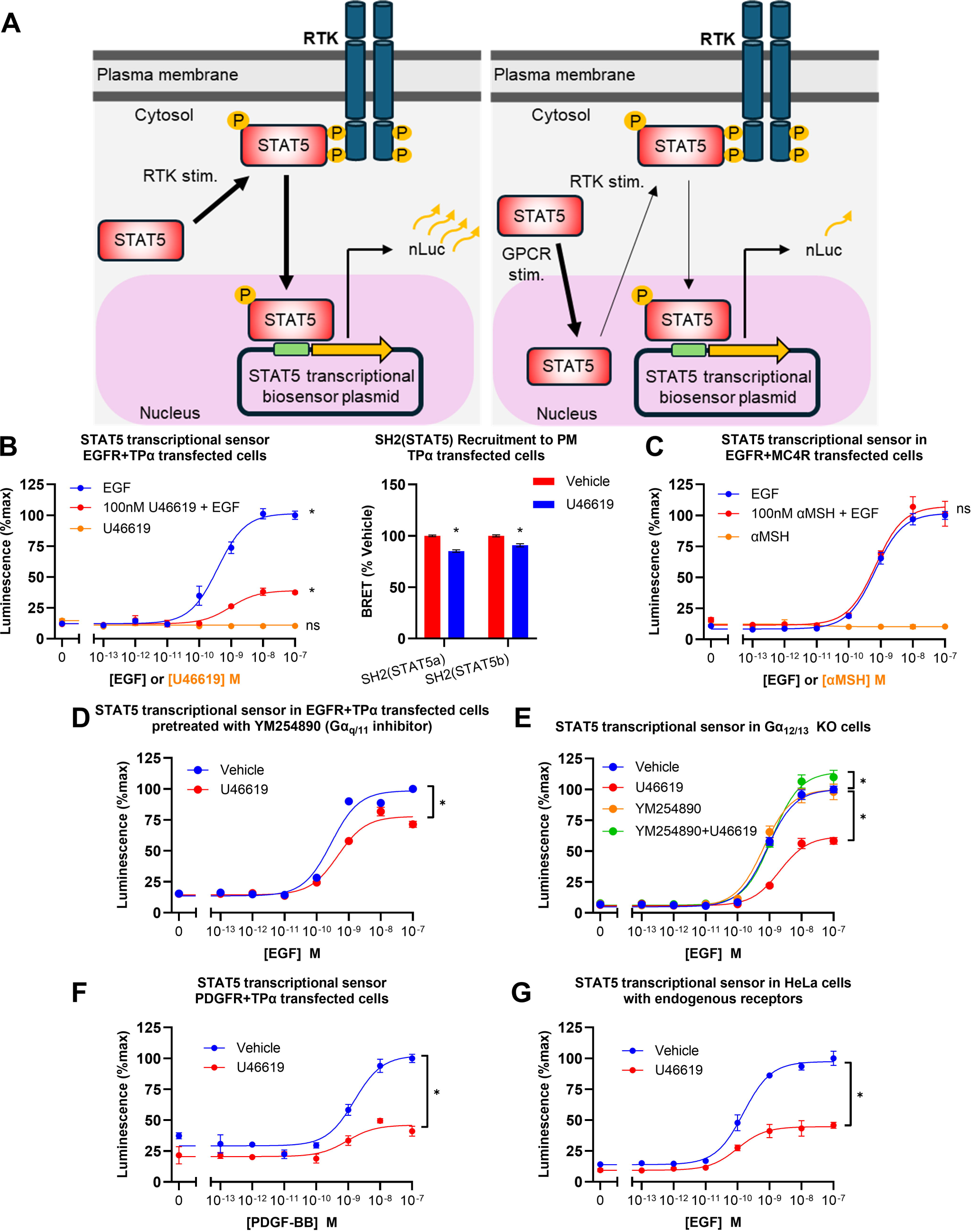
Functional outcomes of GPCR-dependent SH2 domain-containing protein translocation to the nucleus. (A) Schematic representation of STAT5 transcriptional luminescence-based biosensor. This biosensor consists of a plasmid containing an nLuc coding sequence (yellow) under control of a STAT5 specific promoter (green). Activated RTKs recruit endogenous STAT5 through a SH2 domain - phospho-Tyr interaction, resulting in STAT5 phosphorylation and homodimerization. p-STAT5 dimers are translocated to the nucleus and interact with the promoter controlling nLuc expression, allowing for nLuc synthesis and subsequent luminescence. (B, left) HEK293T cells were transfected with the STAT5 transcriptional sensor, EGFR and TPα. Cells were then stimulated with 8 concentrations from 0 to 100nM of EGF (blue line) or 0 to 100nM U46619 (orange line) for 4 hours, or pre-stimulated with a single concentration of 100nM U46619 for 1 hour and treated 8 concentrations from 0 to 100nM of EGF for 4 hours before reading (red line). (B, right) HEK293T cells transfected with TPα, rGFP-CAAX and SH2(STAT5a)-rLuc2 or SH2(STAT5b)-rLuc2, then stimulated with 100nM U46619 for 1 hour before reading. (C) HEK293T cells were transfected with the STAT5 transcriptional sensor, EGFR and MC4R. Cells were then treated with 8 concentrations from 0 to 100nM of EGF (blue line) for 4 hours, or treated with 8 concentrations of αMSH from 0 to 1μM (orange line) for 4 hours or pre-stimulated with a single concentration of 100nM αMSH for 1 hour and treated with 8 concentrations ranging from 0 to 100nM of EGF for 4 hours before reading (red line). (D) HEK293T cells were transfected with STAT5 transcriptional sensor, EGFR and TPα. Cells were then incubated for 30 minutes with 1μM YM254890, pre-stimulated with 100nM U46619 for 1 hour, and treated with 8 concentrations ranging from 0 to 100nM of EGF for 4 hours before reading. (E) HEK293T Gα_12/13_ KO cells were transfected with STAT5 transcriptional sensor, EGFR and TPα. Cells were pre-stimulated with 100nM U46619 for 1 hour and treated with 8 concentrations ranging from 0 to 100nM of EGF for 4 hours before reading in the presence of absence of 1μM Gα_q/11_ inhibitor YM254890. (F) HEK293T cells were transfected with STAT5 transcriptional sensor, PDGFR and TPα. Cells were then pre-stimulated with a 100nM U46619 for 1 hour and treated with 8 concentrations from 0 to 100nM of PDGF-BB for 4 hours before reading. (H) HeLa cells expressing endogenous levels of EGFR and TPα were transfected with STAT5 transcriptional sensor. Cells were then pre-stimulated with a 100nM U46619 for 1 hour and treated with 8 concentrations from 0 to 100nM of EGF for 4 hours before reading. Results are expressed as percentage change relative to GPCR unstimulated control. (mean ± SEM, n=3).

## Discussion

The present study describes a new mechanism by which Gα_q/11_ and Gα_12/13_-coupled GPCRs regulate the activity RTKs. Activation of these G proteins results in a redistribution of SH2 domains found in RTKs effector proteins from the PM to the nucleus, reducing their availability for RTKs engagement (Figure 8) and subsequent signaling. The functional implication of such redistribution was illustrated by the blunted transcriptional activity of STAT5 upon stimulation of either EGFR or PDGFRβ.

**Figure 8.**
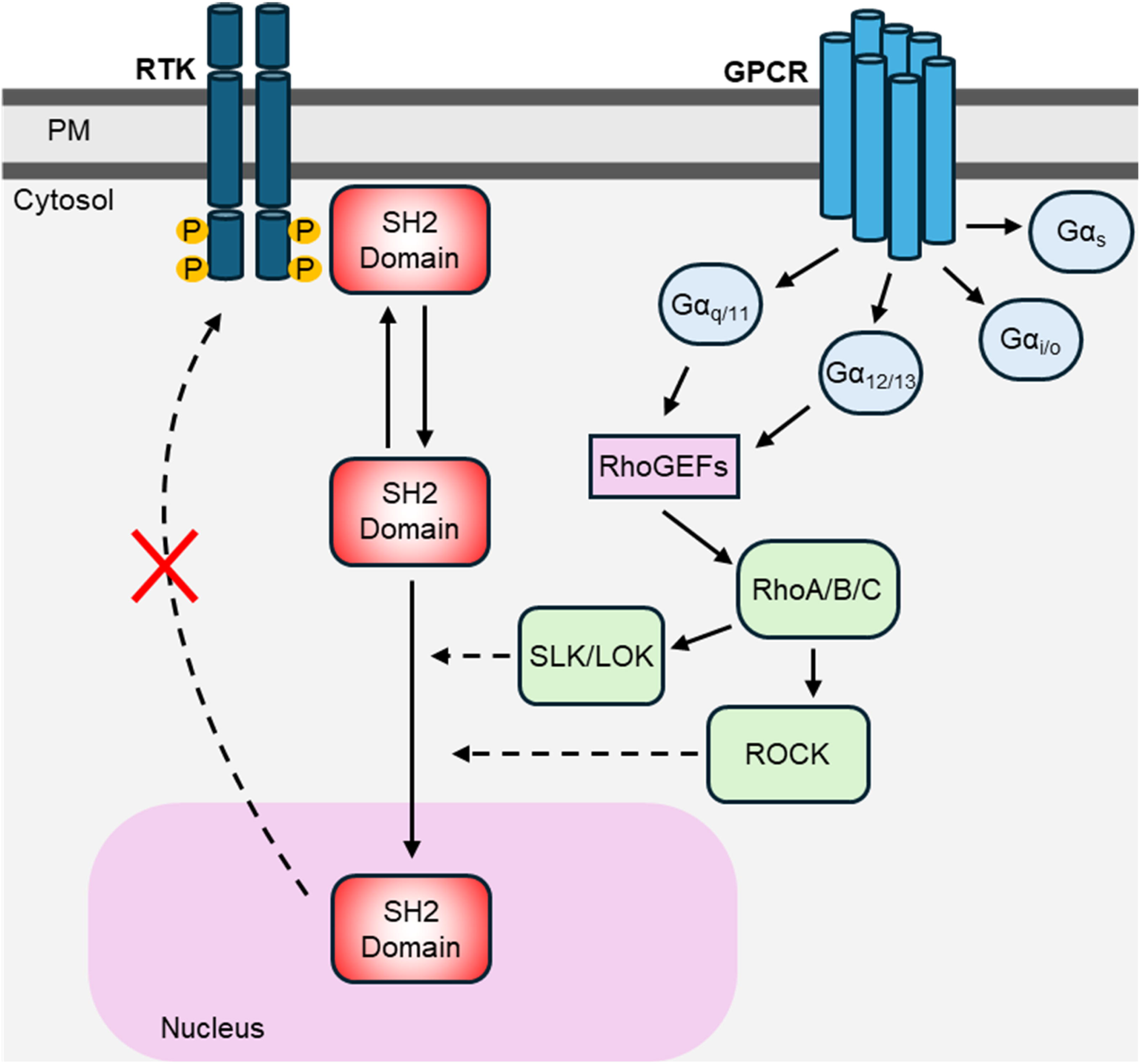
Schematic representation of RhoA-dependent, SH2 domain-containing protein translocation to the nucleus following GPCRs and RTKs activation. The stimulation of GPCRs coupled to Gα_q_ and/or Gα_12/13_ leads to the activation of RhoA/B/C via RhoGEFs. Downstream RhoA/B/C kinases participate in the process of mobilizing SH2 domain-containing proteins away from the plasma membrane and into the nucleus, which can be blocked by CT04, Y-27632 and Cmp31. The compartmentalized SH2 domain-containing proteins are less prone to be recruited by activated RTKs, thus dampening RTKs signal transduction.

RTKs signaling is tightly regulated to prevent pathological overactivation, a hallmark of cancer. RTKs desensitization occurs through a combination of rapid and delayed processes that act in concert to finely regulate receptor activity. Rapid processes are initiated immediately upon activation and include the action of tyrosine phosphatases, receptor ubiquitination for degradation, and the activation of signaling proteins that inhibit specific pathways (30). In contrast, delayed processes are triggered several minutes after receptor activation via negative feedback loops involving proteins such as Sprouty, Sef, and Mig6, which selectively inhibit some RTKs or specific downstream effectors (31). The new regulatory mechanism unravelled here appears to be more general since all tested SH2 domains derived from different effectors dissociated from the PM upon stimulation with Gα_q/11_ or Gα_12/13_-coupled GPCRs.

Cross-activation between GPCRs and RTKs is a widely studied process, with several mechanisms detailed in the literature. Gβγ has been shown to activate matrix metalloproteinases, leading to the release of RTKs ligands and subsequent RTKs activation (32, 33). Also, kinases activated by both GPCRs and RTKs, such as PI3K, Src, and Pyk2, have been shown to phosphorylate RTKs leading to their activation upon GPCRs stimulation (5, 34). Finally, heterodimerization between specific GPCRs and RTKs has been proposed to increase receptor transactivation by forming cooperation hubs consisting of receptors and downstream signaling molecules (35). To date, there are only few examples of cross-inhibition of RTKs activity by GPCRs, including CB1-, CB2- and CXCR4-promoted inhibition of EGFR (11–13). However, the mechanism(s) by which these GPCRs mediate their inhibitory action remain unexplored. CB1, CB2, and CXCR4 are mainly described as Gα_i/o_-coupled receptors. Interestingly, Gα_12/13_ coupling has been described for CB1 (36) and CXCR4 (37), thus suggesting that the previously reported RTK cross-inhibition by these GPCRs may involve the mechanism described herein.

The distinct kinetics of SH2 domain recruitment by RTKs and dissociation from GPCRs suggest different underlying translocation mechanisms. In contrast to the rapid SH2 domain recruitment following RTK stimulation, the slower dissociation observed after GPCR activation likely reflects the involvement of downstream signaling events. This interpretation is consistent with the requirement for Gα_q/11_ / Gα_12/13_ activation and Rho pathway signaling in this process. Additional evidence for the involvement of distinct mechanisms emanates from the phospho-Tyr binding-deficient SH2 domain mutants, which still dissociated from the PM after GPCRs stimulation while failing to respond to classical RTKs activation.

The observation that the phenomenon appears independent of the phospho-Tyr recognition by the SH2 domains raises questions about the mechanism by which these domains associate with the PM at the basal level. Previous studies have suggested interaction motifs beyond phospho-Tyr recognition: 1) surface cationic patches in the SH2 domains can promote association with the internal face of the PM, priming the system for their interaction with RTKs (38); 2) SH2 domains can also bind residues flanking phospho-Tyr, enabling a lower-affinity interaction via their specificity pocket that does not require RTK phosphorylation (39, 40); 3) secondary binding pockets in SH2 domains, such as those formed by the N- and C-terminal regions, promote non-classical interactions with the RTKs (41, 42).

Whether the reduction of SH2 domains from the PM results from a direct action to weaken their spontaneous association with RTKs or the PM or instead reflects a shift in their equilibrium between the PM and the cytosol (potentially influenced by SH2 domain nuclear sequestration) remains to be investigated. The latter hypothesis would suggest that the primary effect would be to facilitate the trafficking of these proteins to the nucleus, a hypothesis supported by our data using the inhibition of importin-β. Although SH2 domain-containing proteins do not possess a canonical nuclear import signal, it is known that many of these proteins (including PLCγ1 (43) and GRB2 (28)) are naturally located inside the nucleus. Therefore, alternative mechanisms possibly involving post-translational modification(s) and/or association with nuclear transport proteins may take place.

We showed that the SH2 domain from STAT5 is translocated to the nucleus after TPα stimulation, and the resulting EGFR signaling through this transcription factor is negatively affected due to its nuclear compartmentalisation. STAT5 was described to be imported to the nucleus independently of RTKs activation following an interaction between its coiled-coil domain and importins through an unconventional NLS signal. However, the function of unphosphorylated STAT5 in the nucleus remains unknown (44). While the translocation of SH2 domain-containing transcription factors such as STAT5 after RTKs activation is a well described process, the mobilization of other SH2 domain-containing proteins to the nucleus is poorly explored. GRB2 can translocate to the nucleus and associate with PTEN and Rad51 to participate in DNA damage repair. In addition, GRB2 participates in double strand break repair by targeting MRE11 nuclease to damage sites through interaction between the GRB2(SH2) and phosphorylated H2AX (28, 45). Nuclear PLCγ1 is an essential part of nuclear inositide signaling cascade, acting together with PI3K and PI3K enhancer (PIKE) to promote cell proliferation and survival (46, 47). Although we examined the effect of GPCR-induced SH2 domain translocation on RTK-mediated transcription, other consequences of GPCR-driven nuclear translocation of SH2 domain-containing proteins may also be functionally significant.

SH2 domain-containing proteins are implicated in various human pathologies beyond cancer, including genetic disorders affecting normal tissue development and immunological disorders (48, 49). Multiple approaches have been undertaken to design molecules that modulate the functionality of SH2 domains related to such conditions. However, the polar nature of key functional regions within this domain (such as the phospho-Tyr-binding and specificity pockets), coupled with a high incidence of off-target effects, presents significant challenges for developing functional compounds capable of crossing the cell membrane. Among the targets investigated to date, notable inhibitors include those directed against PI3K (50), Grb2 (51), STAT3 (52), Zap-70 (53), and Shp2 (54). Our findings suggest that GPCRs-targeted therapies, or interventions disrupting the SH2 domain translocation pathway, could offer new strategies to correct aberrant RTKs signaling, circumventing the problems facing SH2 domain-based drug design.

### Study limitations

A major limitation of the data presented herein is that our BRET-based biosensors consist only of the SH2 domain, and not the full-length protein. Although the STAT5 transcriptional sensor uses endogenous cellular machinery and supports the findings generated with the SH2 domain constructs, the data do not allow us to conclude that all full length SH2 domain-containing proteins would also be imported into the nucleus. Additionally, the roles of SH2 domain-containing proteins translocated into the nucleus following Gα_q/11_ and/or Gα_12/13_ activation is still poorly understood and will be the object of future studies. Finally, while we delineated the GPCRs-Rho-SLK/LOK-ROCK axis as the mechanism acting upstream of SH2 protein redistribution, the effector(s) mediating nuclear translocation remain(s) unknown. Large-scale approaches (e.g., BioID, Co-IP/MS, CRISPR screening) could bridge the gap in the pathway between SLK/LOK-ROK and SH2 domain translocation.

## Materials and Methods

### Cell culture

Human embryonic kidney 293 (HEK293T) cells, human cervical carcinoma cells (HeLa) and human osteosarcoma cells (U2OS) were cultured in complete Dulbecco’s Modified Eagle Medium (DMEM Wisent Inc. #319-015-CL) supplemented with 100 U/mL penicillin and 100 μg/mL streptomycin (Wisent Inc. #450-201-EL) and 10% fetal bovine serum (Wisent Inc. #090150). Cells were incubated at 37°C with 5% CO2 and split twice a week using trypsin (Wisent Inc. #325-542-EL).

### Plasmids

Human SH2 domains from SHP1, SHP2, TNS2, PIK3R2d2, PI3K3R1d1, PLCγ1, GRB2 and GRB14 were synthetized and subcloned in pcDNA3.1(+) containing the rLuc2 upstream of the digestion site for HindIII and XbaI, as previously described (15). SH2(STAT5a) and SH2(STAT5b) sequences were synthetized (Genescript) and subcloned in pcDNA3.1(+) containing the rLuc2 downstream of the digestion site for NheI and AgeI. SH2(PLCγ1)-R37A and SH2(GRB2)-R32A mutants were constructed using the Q5 site directed mutagenesis kit (New England Biolabs) to substitute the arginine by an alanine on the phosphotyrosine binding motif FL(V/I)R using SH2(PLCγ1) and SH2(GRB2) as templates.

The open reading frame for each human receptor (TPα, PAR2, M3R, V2R, AT1R, β2AR, MC4R, A2BR, CXCR4, EGFR, and PDGFR) and for each Gα subunit were cloned in pcDNA3.1(+) expression vector. Both rGFP-CAAX and NLS-rGFP constructs were previously described (16).

### Transfection

DNA mixes were prepared by diluting each DNA to the following concentrations in PBS (concentrations in ng/mL):

For SH2 sensor translocation assays: 10ng of SH2-rLuc2, 300ng of rGFP-CAAX, 50ng of GPCR or 100ng of EGFR. For GPCR activation controls: 10ng of rLuc2 tagged donor (PKN, p63RhoGEF, Rap1Gap, PDZRhoGEF and β-arrestin2), 300ng of rGFP-CAAX, 50ng of GPCR, and 20ng of Gα subunits (Gq for p63RhoGEF, Gi2 for Rap1Gap, G13 for PDZRhoGEF, or Gq, G11, G14, G15, G12, G13 for PKN). For microscopy visualization of BRET: 100ng of SH2-rLuc2, 300ng of rGFP-CAAX or 500ng of NLS-rGFP and 50ng of TPα. For the STAT5 transcriptional biosensor: 100ng of STAT5 biosensor, 100ng of EGFR or PDGFR and 50ng of TPα, PAR2 or MC4R.

The final amount of DNA was corrected for 1μg/mL using salmon sperm DNA as a carrier (Invitrogen, #15632011). The transfection reagent PEI (polyethylenimine 25kDa linear; ThermoFischer Sci. #43896) previously diluted in PBS was added to the DNA mix to reach a final proportion of 3μg PEI to 1μg DNA. DNA:PEI mix was then added to 1,2mL cells in suspension at a density of 3,5×105 cells/mL, mixed and plated in 96 well plates (White Opaque 96-well Microplates; Greiner, # 655083). Plates were then incubated for 48 hours at 37°C with 5% CO2.

### BRET based biosensors

Receptor activation was measured using previously described Effector Membrane Translocation Assay (17) to measure PKN-rLuc2, β-arrestin2-rLuc2, p63RhoGEF-rLuc2 and Rap1GAP-rLuc2 recruitment to the PM resident rGFP-CAAX (16), while EGFR mediated SH2 domain mobilization was measured as described in (15) with minor modifications described below.

48 hours post transfection, cells were washed once with PBS (Wisent Inc. #211410044) and incubated in Tyrode Buffer (137mM NaCl; 0,9mM KCl; 1mM MgCl2; 11,9 NaHCO3; 3,6mM NaH2PO4; 25mM HEPES; 5,5mM D-Glucose; 1mM CaCl2) for at least 1 hour at 37°C. Alternatively, for experiments using Rho, ROCK and SLK/LOK inhibitors (CT-04, Cmpd31 and Y-29632) or experiments using the STAT5 transcriptional sensor, cells were washed once with PBS and incubated in DMEM without FBS, penicillin/streptomycin and phenol red (Wisent Inc. #319051039) for at least 1 hour before treatments.

Cells were stimulated and incubated at 37°C for: 45 minutes for GPCR-mediated translocation of SH2 domain sensors; 10 minutes for EGFR-mediated translocation of SH2-sensors; 10 minutes for GPCR activation controls (PKN, p63RhoGEF, Rap1Gap, PDZRhoGEF, β-arrestin2). 15 minutes before reading, cells were incubated with 1μM methoxy e-Coelenterazine (NanoLight Technology #369).

Data was collected using a SPARK 10M plate reader (Tecan, Switzerland) with the following specifications: 50milisecond integration time; donor filter 400nm ± 40 nm) and acceptor filter (540 ± 35 nm) at 37°C. BRET values were calculated with the ratio of the light emitted by the energy acceptor (rGFP) over the light emitted by the energy donor (rLuc2). For real time kinetics experiments, data was aquired every 12 seconds for 10 minutes before stimulation to stablish the baseline and up to 1 hour after stimulation. Injection was performed using SPARK 10M coupled Injector Module with 200μL/s injection speed.

### Ligands and inhibitors

The following compounds were resuspended / diluted following the manufacturers conditions: Human recombinant Epidermal Growth Factor (EGF) (Sigma Aldrich # 324831); Human PDGF-BB Recombinant Protein (ThermoFischer #100-14B); SLIGKV-NH2 (ApexBio #A8677-5); U46619 (Cayman Chemical #56985-40-1); SQ29,548 (Cayman Chemical #98672-91-4); Carbamoylcholine chloride (Cayman Chemical #51-83-2); Arginine vasopressin (AVP; Sigma-Aldrich #V9879); Isoproterenol hydrochloride (Sigma-Aldrich #I6504); Alpha-MSH (GenScript #RP10644-5); Adenosine (Sigma Aldrich #A9251); Angiotensin II (Sigma-Aldrich # #: A9525); Human Stromal Cell-Derived Factor-1 alpha CXCL12 (Cedarlane #CLY100-20); Rho A/B/C inhibitor, CT04-C3 transferase (Cytoskeleton inc. #CT04); ROCK 1/2 inhibitor, Y-29632 (Sigma Aldrich #688000); SLK/LOK inhibitor, Cmpd31 / SLK/STK10-in-1 (MedChemexpress CO. #HY-132868); Gαq inhibitor, YM-254890 (Cayman Chemical #568580-02-9).

Before data acquisition, each inhibitor was incubated with cells during 6 hours for CT04, 6 hours for Y-29632, 30 minutes for Cmpd31 and 30 minutes for YM-254890.

### Microscopy

Microscopic imaging of BRET signals was performed using photon counting-based method as described in Kobayashi et al. (2019) (18). U2OS cells were seeded on 35mm glass bottom dishes (P35GC-1.5-14-C, MatTek) at a density of 2.0×105 cell per dish and transfected as described in the transfection section. Cells were washed with Modified Hank’s balanced salt solution (HBSS) ((137.9 mM NaCl, 5.33 mM KCl, 1 mM CaCl2, 1 mM MgCl2, 0.44 mM KH2PO4, 0.33 mM Na2HPO4, 10 mM HEPES pH 7.4). Luciferase substrate (Prolume Purple, 10 uM) was diluted with HBSS and added just before the measurement. Microscopic images were obtained with an inverted microscope (Eclipse Ti-E, Nikon, Japan) equipped with x60 objective (CFI Apochromat TIRF, NA 1.49) and separated to donor and acceptor images at 490nm with a dual camera splitter (Twincam, Cairn Research). The beams are introduced into two EMCCD cameras (HNu512, Nuvu Cameras, QC, Canada), and binary photon counting frames were continuously recorded with 100 msec exposure. Final images were obtained by integrating the same numbers of donor frames and acceptor frames until the average photon count of the total luminescence image reaches 100 counts/pixel. BRET image was obtained by dividing acceptor photon counts by donor photon counts, pixel by pixel. To reduce the shot noise level, BM3D filter adapted for Poisson noise reduction (19) was applied and contrast was slightly compressed (gamma = 1.5) for all BRET images. BRET level was described using pseudocolor allocated with ‘jet’ colormap of MATLAB 2021b.

### STAT5 Transcriptional biosensor

To construct an expression vector that measures specifically STAT5 activity, the two following DNA fragments were assembled using Gibson Assembly Cloning (ThermoFischer #A46627): PCR amplified sequence from pUC19 containing AmpR and replication origin; and a synthetic DNA containing polyA sequence on both ends to assure a complete transcriptional isolation of the STAT5 response element, RNA polymerase II transcriptional pause signal from the human α2 globin gene, codon optimized NanoLuc with the 5’ signal peptide from Gaussia Luciferase under control of 5xSTAT5 response element (TTCTGAGA + linker; AAAGTAG) followed by TATA-box minimal promoter.

Following transfection and cell washing as previously described, cells transfected with STAT5 transcriptional biosensor were stimulated with increasing concentrations of EGF or increasing concentrations of U46619 and incubated in DMEM without FBS, penicillin/streptomycin and phenol red for 4 hours at 37°C with 5% CO2. For experiments with GPCR pre stimulation, cells were stimulated with U46619, αMSH or SLGKV-NH2 for 1 hour prior EGF addition. For experiments with both GPCR pre stimulation and Gαq inhibition, YM-254890 was added 30 minutes before U46619 addition. After incubation with agonists, Furimazine (Molnova #M23530) was added to each well to a final concentration of 1μM and proceeded to reading immediately on SPARK 10M plate reader. The parameters were as follows: total luminescence reading with integration time of 200 milliseconds at 37°C.

## Acknowledgments

The authors are grateful to Monique Lagacé for her help in the preparation of the manuscript. The work was supported by a CIHR project grant (PJT-183758) and NSRC discovery grant to MB. PHSP held a MITACS Accelerate fellowship.

## Supplementary Figure Legends

**Figure S1.** Gα protein dependent SH2(PLCy1) mobilization by PAR2. HEK293T Gα Knockout cells were transfected with SH2(PLCγ1)-rLuc2, rGFP-CAAX and PAR2. Gα protein expression was rescued by transfecting 0, 5, 20 or 50ng/μg DNA of Gα_q_ (A), Gα_i2_ (B) or Gα_13_ (C). Cells were treated with SLIGKV-NH2 and signals were read after 45 minutes to calculate BRET values. As a control for Gα protein rescue, HEK293T Gα Knockout cells were transfected with PAR2, rGFP-CAAX and specific sensor for each Gα Family (p63RhoGEF-rLuc2 for Gα_q_ (D), Rap1Gap-rLuc2 for Gα_i2_ (E) and PDZRhoGEF-rLuc2 for Gα_13_ (F)). Cells were treated with SLIGKV-NH2 and signals were read after 1 minute. BRET values were calculated by dividing the intensity of light emitted by rGFP (515 nm) by the intensity of light emitted by rLuc2 (400 nm). Results are expressed as percentage change relative to unstimulated control (mean ± SEM, n=3). * Statistically significant difference by two-way ANOVA with Šidák post-test, p<0,001.

**Figure S2.** Schematic representation of STAT5 transcriptional biosensor plasmid. From left to right: poly(A) signal sequence + pause site were added to block and transcription of AmpR gene due to minimal promoter activity; STAT5 response elements contains 5x repetitions of binding motifs; the minimal promoter + Kozak sequence allows the correct positioning for gene transcription after STAT5 binding; the signal peptide (blue) was added to allow Nluc export to the extracellular milieu to allow measurements of luminescence using cells supernatant; Nluc gene (purple) followed by a second poly(A) signal sequence to allow transcription end; origin or replication (yellow) and AmpR gene (green).

## Notes

### Competing Interest Statement

MB is a member of the scientific advisory board of Kainova Therapeutics
AM, LS are paid employees of Kainova Therapeutics

