## Supplementary figures and images for "Unconventional Interplay Between GPCRs and RTKs Signaling Pathways Through SH2 Domain-Containing Proteins"

### Figure S1.tif

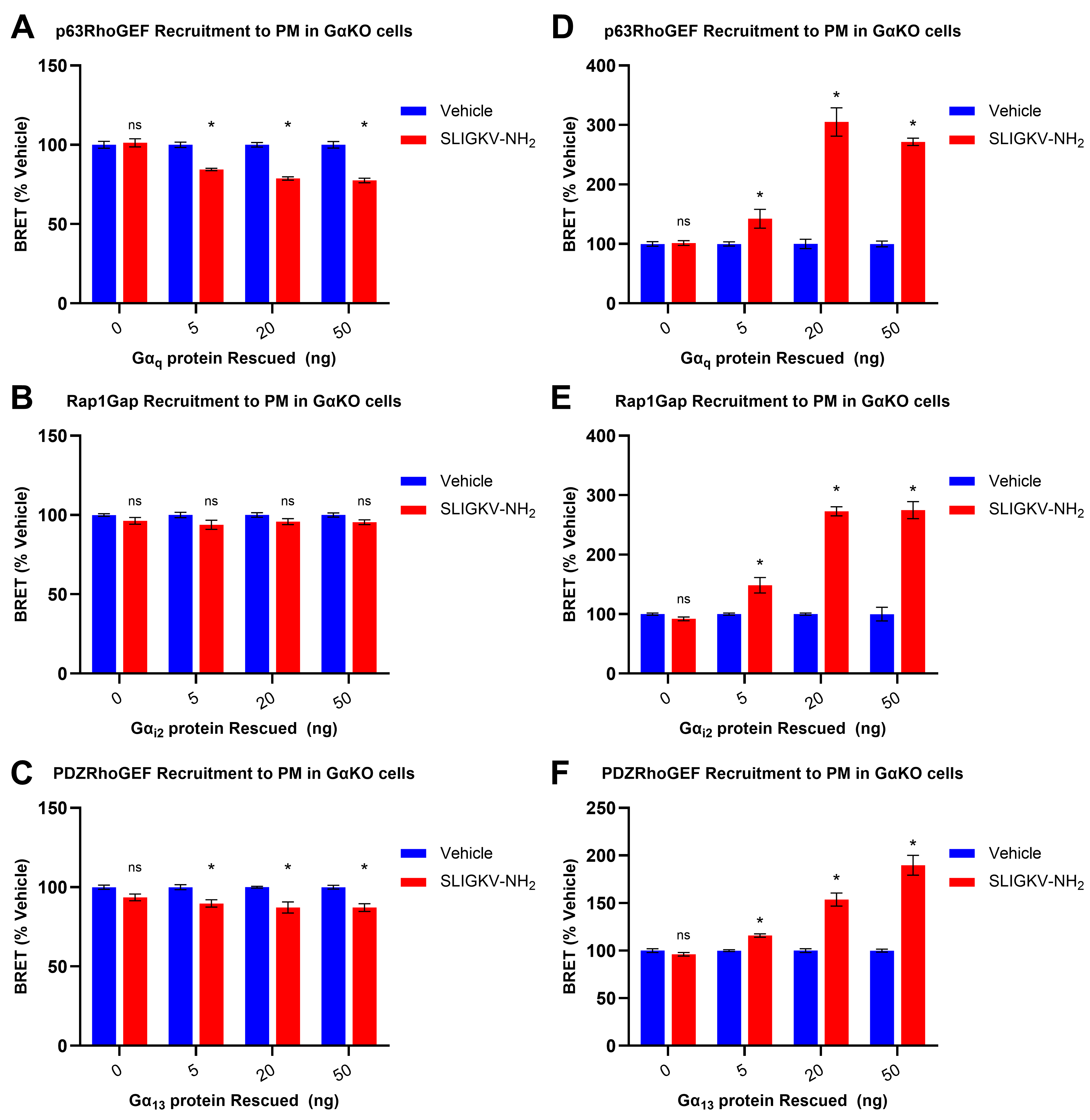

### Figure S2.tif

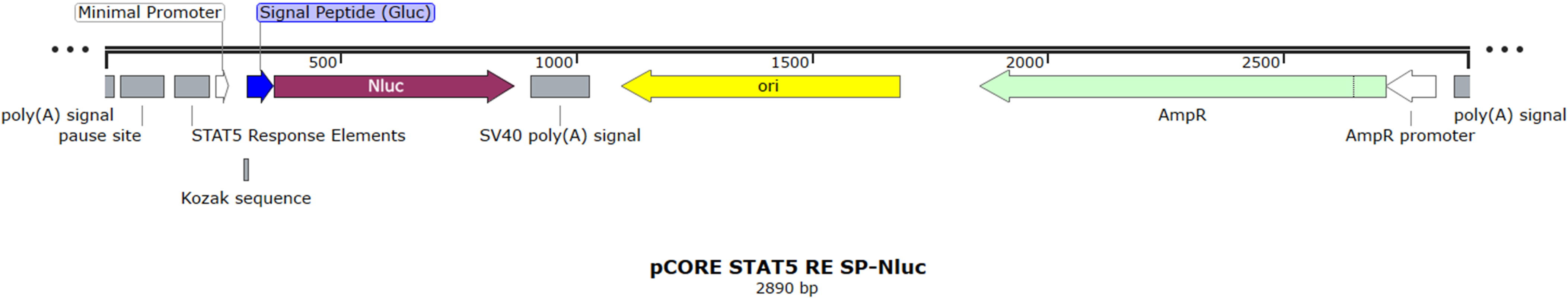
